# EcoFoldDB: Protein structure-guided functional profiling of ecologically relevant microbial traits at the metagenome scale

**DOI:** 10.1101/2025.04.02.646905

**Authors:** Timothy M. Ghaly, Vaheesan Rajabal, Dylan Russell, Elena Colombi, Sasha G. Tetu

## Abstract

Microbial communities are fundamental to planetary health and ecosystem processes. High-throughput metagenomic sequencing has provided unprecedented insights into the structure and function of these communities. However, functionally profiling metagenomes remains constrained due to the limited sensitivity of existing sequence homology-based methods to annotate evolutionarily divergent genes. Protein structure, more conserved than sequence, and intrinsically tied to molecular function, offers a solution. Capitalising on recent breakthroughs in structural bioinformatics, we present EcoFoldDB, a database of protein structures curated for ecologically relevant microbial traits, and its companion pipeline, *EcoFoldDB-annotate*, which leverages Foldseek with the ProstT5 protein language model for rapid structural homology searching directly from sequence data. *EcoFoldDB-annotate* outperforms state-of-the-art sequence-based methods in annotating metagenomic proteins, in terms of sensitivity and precision. To demonstrate its utility and scalability, we performed structure-guided functional profiling of 32 million proteins encoded by 8,000 high-quality metagenome-assembled genomes from the global soil microbiome. *EcoFoldDB-annotate* could resolve the phylogenetic partitioning of important nitrogen cycling pathways, from taxonomically restricted nitrifiers to more widespread denitrifiers, as well as identifying novel, uncultivated bacterial taxa enriched in plant growth-promoting traits. We anticipate that EcoFoldDB will enable researchers to extract ecological insights from environmental genomes and metagenomes, and accelerate discoveries in microbial ecology.

## Introduction

Microbial community functioning shapes Earth system health, influencing global carbon dynamics, driving essential nutrient cycles and ecosystem services, and underpinning the biodiversity of entire biomes (Wardle et al., 2004; Gillings and Paulsen, 2014; Steffen et al., 2015; Cavicchioli et al., 2019). Deciphering the complexity of microbial communities, including their composition, interactions, and functions, is critical for understanding ecosystem processes and their responses to environmental change. High-throughput shotgun metagenomic sequencing has revolutionised the field of microbial ecology, allowing a culture-independent view of the structure and function of microbiomes (Huttenhower et al., 2012; Sunagawa et al., 2015; Bahram et al., 2018; Bahram et al., 2022), and facilitating unprecedented insights into their ecology and evolution at scales ranging from individual strains to whole communities (Rohwer et al., 2025; Zhou et al., 2025).

Functional annotation of microbial genes represents a cornerstone of metagenomic analysis. Nevertheless, this task represents a formidable challenge due to the extensive size of metagenomic datasets, and the massive functional diversity encoded within metagenomes (Pavlopoulos et al., 2023). Several bioinformatic tools exist to enable large-scale functional profiling, e.g., InterProScan (Jones et al., 2014), eggNOG-mapper (Cantalapiedra et al., 2021), KofamScan (Aramaki et al., 2019). However, these tools prioritise exhaustive functional identification over ecological interpretation. Researchers thus often need to manually screen annotation results to gain meaningful insights into the ecological functions of microbial communities. This has led to the development of ecology-focused tools for functional profiling, such as METABOLIC (Zhou et al., 2022) and DRAM (Shaffer et al., 2020), which curate annotations into ecologically coherent functional categories, enabling user-friendly interpretation and analysis. However, existing functional annotation tools rely on sequence homology-based methods (e.g., MMSeqs2 (Steinegger and Söding, 2017), DIAMOND (Buchfink et al., 2021), or HMMER (Finn et al., 2011)). Consequently, annotation of metagenomes remains constrained since they are overwhelmingly composed of genes from uncultivated microbial taxa, and are enriched with proteins lacking detectable sequence homology to reference databases (Vanni et al., 2022; Pavlopoulos et al., 2023).

The structure of proteins, which are inherently linked to their molecular function, allows homology discovery beyond the limits of sequence similarity detection, as structure evolves much slower than sequence (Illergård et al., 2009). However, until recent breakthroughs in structural bioinformatics, the use of protein structure for large-scale functional profiling has not been feasible. Notably, AlphaFold2 revolutionised protein structure predictions, achieving near-experimental accuracy from sequence data alone (Jumper et al., 2021). Yet, identifying similar protein structures at scale remained a significant challenge. Foldseek addressed this limitation by enabling protein structural homology searching at speeds comparable to sequence searching (van Kempen et al., 2024). Foldseek achieves this by mapping the 3D coordinates for each residue in a protein structure to one of the 20 states in their 3D interaction (3Di) alphabet. Thus, Foldseek effectively represents the structure of a protein as a 20-letter alphabet, enabling the use of fast sequence searching algorithms that were originally developed for the 20-letter amino acid alphabet (Farrar, 2007; Zhao et al., 2013; Steinegger and Söding, 2017). However, despite these advances, the computational bottleneck of AlphaFold2 structure predictions still hindered metagenomic applications, even with performance optimisations like ColabFold (Mirdita et al., 2022). To that end, the ProstT5 protein language model was developed, which translates between protein amino acid and Foldseek’s 3Di sequences, without the need for structure predictions (Heinzinger et al., 2024). ProstT5 was built from an existing metagenomic protein language model (ProtT5) (Elnaggar et al., 2021), and further trained on 17 million non-redundant and diverse protein sequences, paired with their corresponding 3Di sequences from the AlphaFold Protein Structure Database (Varadi et al., 2023). Consequently, ProstT5 enables ∼4,000× faster structural homology searching, compared to AlphaFold2-ColabFold, without sensitivity loss (Heinzinger et al., 2024).

To take advantage of these recent breakthroughs for applications in microbial ecology, we created EcoFoldDB, a curated Foldseek database of protein structures spanning microbial traits of ecological relevance. The included traits cover biogeochemical cycling of carbon, nitrogen, sulphur, phosphorus, iron, and trace atmospheric gases, as well as key plant-microbe interactions, and osmotic stress tolerance mechanisms. Complementing this resource, we introduce *EcoFoldDB-annotate*, an annotation pipeline that leverages the speed and accuracy of ProstT5’s 3Di sequence translation, and Foldseek’s structural homology search capabilities. This annotation framework enables structure-guided functional profiling, scalable to metagenomic datasets.

## Methods

### Database construction

We included eight microbial functional categories, and 35 sub-categories, within EcoFoldDB that span several key ecological processes (Supplementary Table S1). Pathways and associated genes were sourced from published literature or existing functional databases.

These consisted of *(1) Trace gas oxidation*: atmospheric hydrogen oxidation (Søndergaard et al., 2016; Bay et al., 2021), carbon monoxide oxidation (Bay et al., 2021), and methane oxidation (Bay et al., 2021); *(2) Carbon cycling*: carbon fixation (Caspi et al., 2013; Ruiz-Fernández et al., 2020; Garritano et al., 2022), one-carbon (C1) metabolism (Caspi et al., 2013), complex carbohydrate degradation (Zheng et al., 2023), fatty acid degradation (Caspi et al., 2013), and polyphenol and aromatic hydrocarbon cycling (McGivern et al., 2024b; McGivern et al., 2024a); *(3) Nitrogen cycling*: nitrogen fixation, nitrification, denitrification, dissimilatory nitrate reduction to ammonium (DNRA), nitrate assimilation, anaerobic ammonium oxidation (anammox), and organic nitrogen mineralisation (Caspi et al., 2013; Tu et al., 2018; Ghaly et al., 2025); *(4) Sulphur cycling*: sulphate reduction, sulphite reduction, sulphide oxidation, sulphur disproportionation, sulphite oxidation, thiosulphate oxidation, thiosulphate disproportionation, sulphur reduction, and organic sulphur mineralisation (Morra and Dick, 1991; Eichhorn et al., 1997; Soutourina et al., 2001; Jamshad et al., 2007; Ellis, 2011; Caspi et al., 2013; Zhou et al., 2022); *(5) Phosphorus cycling*: organic phosphorus mineralisation and inorganic phosphorus solubilisation (Siles et al., 2022; Zeng et al., 2022; Ghaly et al., 2025); *(6) Iron cycling*: siderophore biosynthesis (Caspi et al., 2013); *(7) Plant-microbe interactions*: ethylene suppression via ACC deaminase activity (Glick et al., 2007), indole-3-acetic acid production (Tang et al., 2023), gibberellin production (Bottini et al., 2004), abscisic acid production (Khalil et al., 2024), trans-Zeatin production (Frébortová and Frébort, 2021), jasmonic acid production (Li et al., 2005; Khalil et al., 2024), brassinolide production (Bajguz et al., 2020), GABA production (Kinnersley and Turano, 2000; Ueno, 2000; Ma et al., 2012), putrescine and spermidine production (Dunn and Becerra-Rivera, 2023), and legume root nodulation (Rossen et al., 1985; Prell and Poole, 2006); and (8) *Osmotic stress tolerance*: osmoprotectant biosynthesis (Jebbar et al., 1992; Chen and Beattie, 2008; Ruhal et al., 2013; Yang et al., 2024). Included genes and pathways, and their functional categorisations are available as Supplementary Table S1.

Proteins with these functional annotations were selected from the manually curated databases, Swiss-Prot (Boeckmann et al., 2003) and MetaCyc (Caspi et al., 2013). The corresponding predicted structures for these protein sequences were then retrieved from the AlphaFold Protein Structure Database (AFDB) (Jumper et al., 2021; Varadi et al., 2023), with protein fragments discarded. Redundancy was removed from this set of protein structures using Foldseek *cluster* (Barrio-Hernandez et al., 2023; van Kempen et al., 2024) with a minimum TM-score threshold of 0.9 and a bidirectional coverage of 0.9 [parameters: foldseek cluster --alignment-type 1 --tmscore-threshold 0.9 -c 0.9 --cov-mode 0 --cluster-reassign 1]. This final set of target EcoFoldDB structures were combined with structures of non-target proteins (i.e., those not included in our curated set of functions) from AFDB-Swiss-Prot. This was done to reduce false positives, by allowing the distinction between structurally homologous proteins with divergent functions.

### EcoFoldDB-annotate pipeline and performance benchmarking

To enable metagenome-scale annotations using EcoFoldDB, we created *EcoFoldDB-annotate*, which employs Foldseek (van Kempen et al., 2024) with the ProstT5 protein language model (Heinzinger et al., 2024). Together, Foldseek with ProstT5, allow protein structural alignments from amino acid sequences without the need for structure predictions (Heinzinger et al., 2024). *EcoFoldDB-annotate* begins by filtering protein sequences exceeding 4,000 amino acids in length, archiving excluded sequences in a dedicated output file for user inspection. Proteins larger than this drastically increase the runtime of ProstT5 inference, and are likely too large to represent any protein in EcoFoldDB, with its largest entry being 2,378 amino acids. *EcoFoldDB-annotate* then uses ProstT5 to translate the amino acid alphabet to Foldseek’s 3D interaction (3Di) structural alphabet, allowing structural alignment of input proteins against EcoFoldDB with Foldseek. Finally, the pipeline assigns functionally categorised annotations based on EcoFoldDB ontology, using the best-hit approach with default query and target coverage cut-offs of 80%, and an e-value threshold of 1e-10.

To test its functional annotation capacity, we benchmarked *EcoFoldDB-annotate* against two widely used state-of-the-art annotation tools, eggNOG-mapper v2.1.13 (Cantalapiedra et al., 2021) and KofamScan v1.3.0 (Aramaki et al., 2019). To create a benchmarking dataset with ‘known’ positives, we used the ESM Metagenomic Atlas 30 (Lin et al., 2023), a non-redundant database of 37 million high-confidence (pTM > 0.7 and pLDDT > 0.7) predicted protein structures from metagenomic assemblies (derived from the MGnify database (Richardson et al., 2022)). The ESM Atlas 30 was chosen as it includes diverse proteins found in metagenomes, with protein entries sharing <30% sequence similarity with each other, and includes protein ‘dark matter’ sequences from genomes of uncultivated taxa. To identify a set of ground truth proteins, we first generated a subset of ESM Atlas proteins exhibiting high structural similarity to those encoding target functions, as determined by Foldseek. We selected only hits with both a query and target TM-score > 0.8, since this threshold indicates a near certain (∼100%) posterior probability of sharing the same fold topology (Xu and Zhang, 2010). Next, the functions of these structural homologs were predicted using CLEAN v1.0.1 (Yu et al., 2023), which uses a contrastive machine learning model for high-quality enzymatic function predictions. Thus, in the absence of a validated reference dataset, these high-confidence matches (i.e., from both structural homology and CLEAN enzyme predictions) were used as a proxy for ground truth functional annotations.

This resulted in 36,639 proteins to serve as ‘known’ positives. To include known negatives in the benchmarking dataset, we used all Swiss-Prot entries with non-target functions based on their assigned EC (Enzyme Commission) numbers. We limited this to Swiss-Prot entries (as of July 2025) not in AFDB-Swiss-Prot (derived from UniProt April 2021 data) as the latter are included in EcoFoldDB. This resulted in 4,106 proteins to act as known negatives.

The corresponding amino acid sequences of the combined positive and negative benchmarking dataset were then used as input for *EcoFoldDB-annotate*, eggNOG-mapper, and KofamScan. *EcoFoldDB-annotate* was run with default settings; eggNOG-mapper sequence searches were performed using DIAMOND (Buchfink et al., 2021) against eggNOG v5.0.2 orthology data (Huerta-Cepas et al., 2018); and KofamScan was run using its model-specific score thresholds (Aramaki et al., 2019).

### EcoFoldDB annotations of microbial genomes from the global soil microbiome

To demonstrate its utility and scalability, we employed *EcoFoldDB-annotate* on >8,000 high-quality metagenome-assembled genomes (MAGs) from the global soil microbiome. Genome sequences were obtained from the Soil Microbial Dark Matter MAG (Ma et al., 2023) and Microflora Danica long-read MAG (Sereika et al., 2024) catalogues, collectively covering diverse soil habitats sampled across the globe. Genome sequences were filtered to retain only high-quality (HQ) MAGs based on MIMAG standards (Bowers et al., 2017). That is, genomes that are >90% complete, with <5% contamination, have the 23S, 16S, and 5S rRNA genes, and at least 18 different tRNA genes. Genome completion and contamination was determined for all MAGs using CheckM2 (Chklovski et al., 2023) with default settings.

Genomes were screened for rRNA genes using barrnap (https://github.com/tseemann/barrnap) and bakta v1.8.1 (Schwengers et al., 2021) (which uses Infernal (Nawrocki and Eddy, 2013) against the Rfam database (Ontiveros-Palacios et al., 2025)), and tRNA genes were screened using bakta (which employs tRNAscan-SE 2.0 (Chan et al., 2021)) with default settings. MAGs were dereplicated at approximately species-level (95% average nucleotide identity, ANI, and 50% aligned fraction, AF) with dRep v3.5.0 (Olm et al., 2017). dRep was run using only skani v0.2.2 (Shaw and Yu, 2023) for ANI and AF calculations, and CheckM2 results were provided to dRep for genome quality information [command: dRep dereplicate --SkipMash --S_algorithm skani -sa 0.95 -nc 0.5 --genomeInfo checkm2_info.csv -p 48].

Proteins were predicted from the HQ soil MAGs using Prodigal v2.6.3 (Hyatt et al., 2010) [parameters: -p meta], which was parallelised across 48 CPU cores using GNU Parallel (Tange, 2018). This resulted in the prediction of 31,890,584 proteins. These proteins were functionally annotated against EcoFoldDB using *EcoFoldDB-annotate* with default settings. Genome sequences and predicted protein sequences of these HQ MAGs have been made available via https://doi.org/10.5281/zenodo.15054612, and EcoFoldDB annotations are available as Supplementary Table S2.

Taxonomy of the HQ soil MAGs was assigned with GTDB-Tk v2.4.0 (Chaumeil et al., 2019, 2022) using GTDB release 220 (Parks et al., 2018; Parks et al., 2020; Parks et al., 2021; Rinke et al., 2021) [parameters: classify_wf --skip_ani_screen]. A whole-genome phylogeny was generated using PhyloPhlAn 3.0 (Asnicar et al., 2020), employing DIAMOND (Buchfink et al., 2021) for marker detection, MAFFT (Katoh and Standley, 2013) for sequence alignment, trimAl (Capella-Gutiérrez et al., 2009) for alignment trimming, and FastTree2 (Price et al., 2010), with the WAG substitution model (Whelan and Goldman, 2001), for phylogenetic inference [PhyloPhlAn parameters: --diversity high --fast]. The resulting phylogenetic tree was rooted with minimal ancestor deviation (MAD) rooting (Tria et al., 2017) using MADroot (Bryant and Charleston, 2018) (https://github.com/davidjamesbryant/MADroot), and visualised using the *ggtree* (Yu et al., 2017) and *ggtreeExtra* (Xu et al., 2021) R packages.

## Results and Discussion

Here, we present EcoFoldDB, a database of protein structures that cover microbial functions of ecological relevance, including those involved in: (1) trace gas oxidation, (2) carbon cycling, (3) nitrogen cycling, (4) sulphur cycling, (5) phosphorus cycling, (6) iron cycling, (7) plant-microbe interactions, and (8) osmotic stress tolerance (Fig. 1). These eight functional categories are organised into 35 sub-categories, and encompass 166 different functional activities/pathways (Fig. 1; Supplementary Table S1). Together, EcoFoldDB comprises 842 protein structures linked to 637 genes (Supplementary Table S1). These functions cover key biogeochemical cycles important for ecosystem health, biodiversity, and climate change (i.e., carbon sequestration and greenhouse gas emissions), as well as microbial traits relevant to adaptive responses to environmental stress, and plant growth promotion (Fig. 1). Future versions of EcoFoldDB can be readily expanded to incorporate new functions, particularly as novel microbial traits influencing ecosystem dynamics are uncovered.

**Fig. 1.**
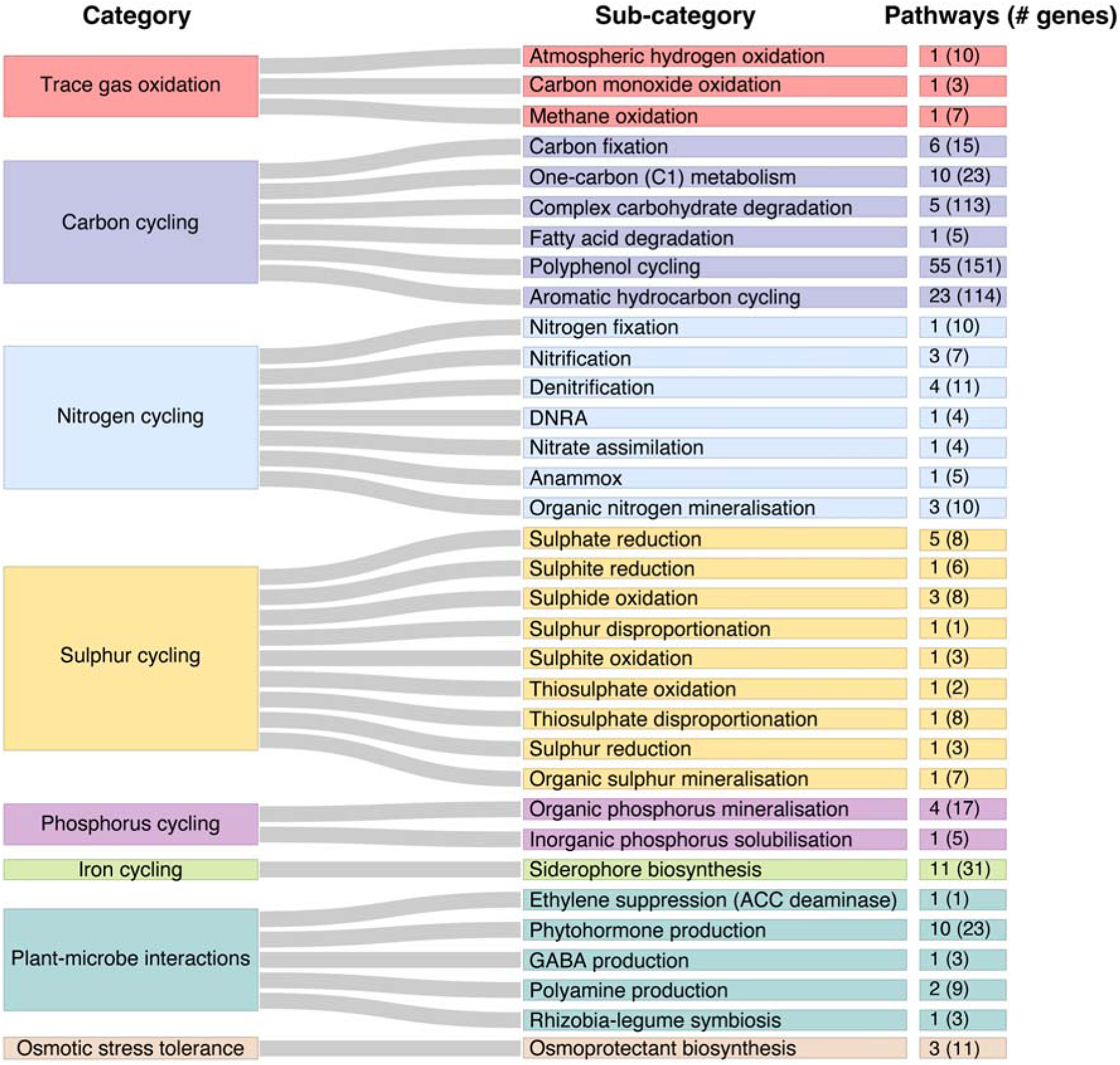
Ecologically relevant functions covered by EcoFoldDB. The EcoFoldDB ontology is comprised of 8 functional categories (left), which are divided into 34 sub-categories (middle). These cover 166 functional pathways/activities and 637 gene families (right). These gene families are represented by 842 protein structures in EcoFoldDB. For a full list of EcoFoldDB entries and their functional categorisations, see Supplementary Table S1.

### ‘EcoFoldDB-annotate’ performs highly sensitive and accurate structure-guided functional profiling

To enable scalable, structure-guided functional annotation using EcoFoldDB, we developed *EcoFoldDB-annotate* (Fig. 2a). This pipeline leverages Foldseek (van Kempen et al., 2024) and the ProstT5 protein language model (Heinzinger et al., 2024) to directly translate amino acid sequences into 3Di structural representations, enabling homology searches against EcoFoldDB without the need for computationally intensive structure predictions. ProstT5 achieves this translation ∼4,000-fold faster than AlphaFold2-ColabFold without losing sensitivity (Mirdita et al., 2022), allowing structural homology detection scalable to metagenomic datasets.

**Fig. 2.**
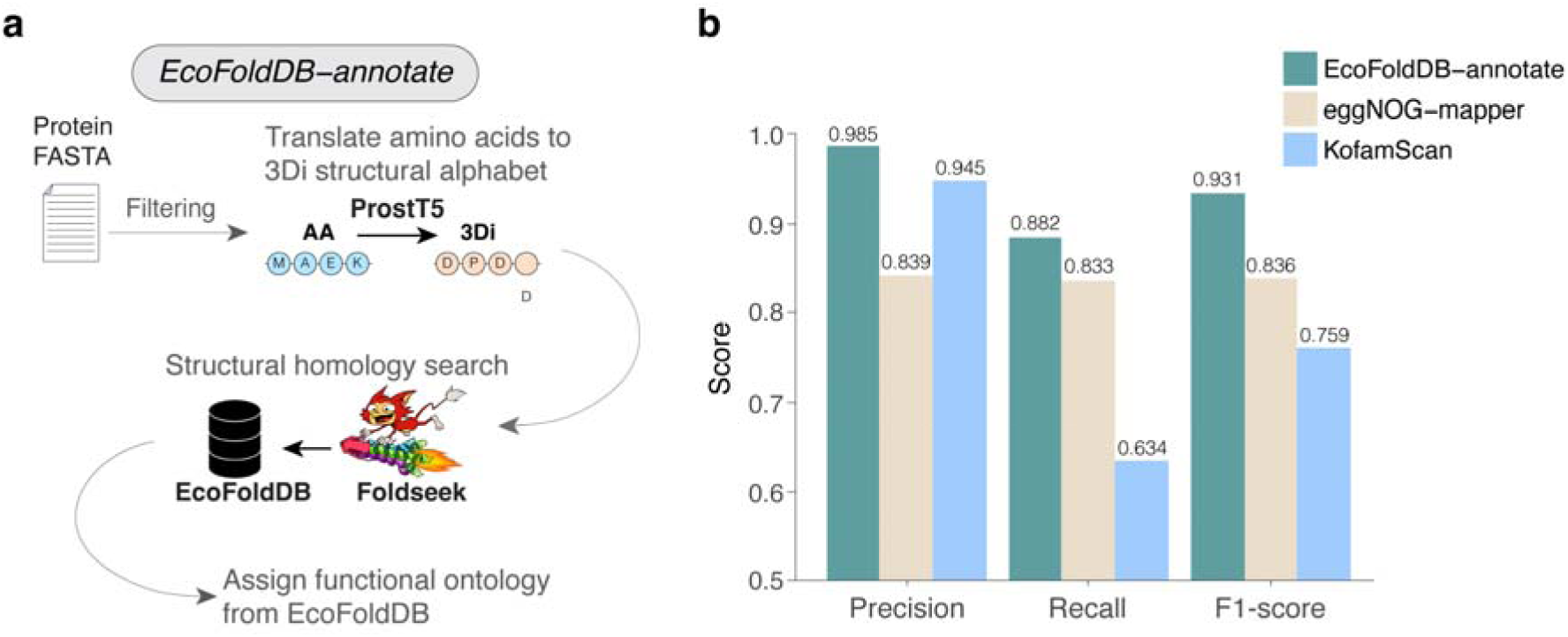
*EcoFoldDB-annotate* pipeline and performance benchmarking. **a**, *EcoFoldDB-annotate* first filters protein sequences to remove those larger than 4,000 amino acids in length, then uses the ProstT5 protein language model to translate amino acid sequences to Foldseek’s 3D interaction (3Di) structural alphabet for structural homology searching against EcoFoldDB, and finally assigns hierarchically categorised functional annotations based on EcoFoldDB ontology. **b**, The performance of *EcoFoldDB-annotate* was benchmarked against eggNOG-mapper and KofamScan. The following benchmarking metrics were used: recall/sensitivity (proportion of all possible positives correctly identified), precision (proportion of predicted positives that were true), and F1-score (the harmonic mean of recall and precision).

*EcoFoldDB-annotate* outperforms sequence-based annotations for proteins derived from metagenomic assemblies (Fig. 2b). *EcoFoldDB-annotate* achieves higher sensitivity, with a recall of 0.882 compared to 0.833 (eggNOG-mapper) and 0.634 (KofamScan), while maintaining the highest precision (0.985) and F1-score (0.931) among the three tools (Fig. 2b). This combination of high sensitivity and precision establishes *EcoFoldDB-annotate* as an effective solution for annotating metagenomes, which are frequently dominated by sequences lacking detectable amino acid homology to reference databases (Vanni et al., 2022; Pavlopoulos et al., 2023). Its performance, scalability, and hierarchical classification of ecologically relevant functions (Fig. 1) positions *EcoFoldDB-annotate* as an invaluable resource for environmental metagenomic studies.

### EcoFoldDB-annotate is scalable to millions of proteins from soil metagenomes

To demonstrate the utility of *EcoFoldDB-annotate*, we used it to perform structure-based functional annotations of 32 million protein sequences encoded by 8,000 high-quality metagenome-assembled genomes (MAGs), derived from diverse soil habitats across the globe. Soil microbes harbor a diverse range of metabolic capabilities, and influence global carbon dynamics and biogeochemical cycles that shape ecosystem health and productivity (Fierer, 2017). Soil environments generally harbour greater phylogenetic diversity and genetic novelty compared to other environments (Thompson et al., 2017; Brewer et al., 2019; Holland-Moritz et al., 2021), making sequenced-based functional annotations particularly difficult (Holland-Moritz et al., 2021). Leveraging the recent breakthroughs in protein structure predictions (Jumper et al., 2021; Mirdita et al., 2022) can improve functional annotations of proteins that lie beyond the limits of sequence homology detection with known proteins (Durairaj et al., 2023). To that end, we employed *EcoFoldDB-annotate* to functionally annotate MAGs from the global soil microbiome. We processed 32 million proteins in batches of 1 million sequences, taking ∼5.3 hours per batch using a server with 4 Nvidia Tesla Volta V100 GPUs and 48 CPU cores (see Supplementary Table S2 for all EcoFoldDB annotations). This demonstrates the scalability of *EcoFoldDB-annotate* to functionally profile large metagenomic datasets.

### EcoFoldDB reveals the taxonomic distribution of nitrogen cycling functions in the global soil microbiome

As a use case example, we used the EcoFoldDB annotations of the high-quality MAGs to reveal the phylogenetic distribution of encoded nitrogen cycling pathways in soils. Our findings reveal distinct taxonomic partitioning, with some pathways, particularly nitrification, being taxonomically restricted, while others, like dissimilatory nitrate reduction (DNRA) and denitrification, were broadly distributed across taxa (Fig. 3). For instance, ammonia oxidation, the initial step in nitrification, was restricted to just three prokaryotic families.

**Fig. 3.**
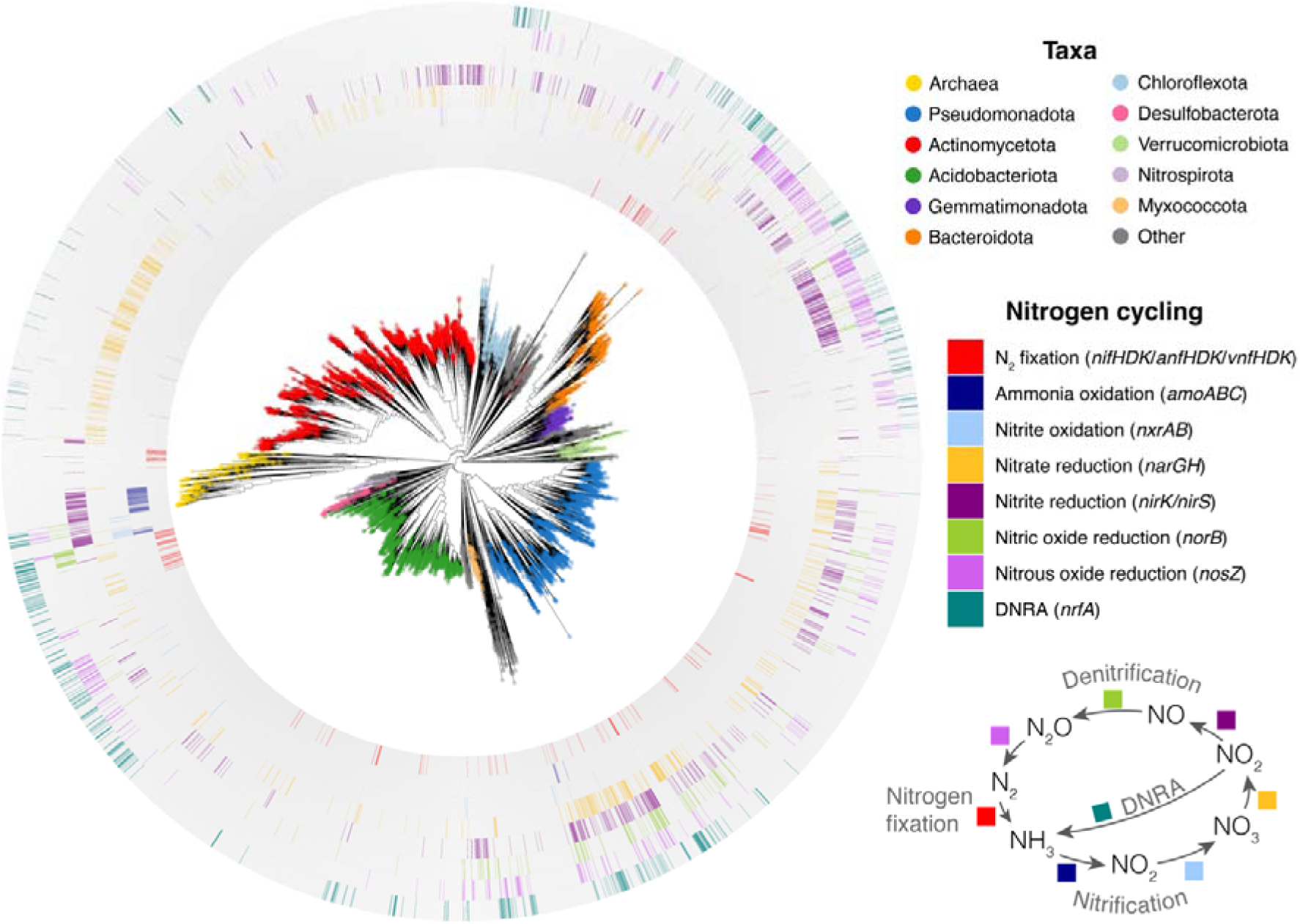
Nitrogen cycling functions identified by *EcoFoldDB-annotate* among high-quality MAGs from the global soil microbiome. A whole genome phylogeny of high-quality bacterial and archaeal MAGs, derived from soil metagenomes across the globe. Coloured tip points on the tree indicate GTDB phyla, except for archaeal genomes, which have all been assigned a single domain-level colour (yellow tip points). Coloured rings indicate the genomes identified to have the genetic potential for nitrogen cycling functions. As a minimum, the following genes were required for N_2_ fixation: (*nifH* and *nifD* and *nifK*) or (*anfH* and *anfD* and *anfK*) or (*vnfH* and *vnfD* and *vnfK*); ammonia oxidation: *amoA*, *amoB* and *amoC*; nitrite oxidation: *nxrA* and *nxrB*; nitrate reduction: *narG* and *narH*; nitrite reduction: *nirK* or *nirS*; nitric oxide reduction: *norB*; nitrous oxide reduction: *nosZ*; and DNRA (dissimilatory nitrite reduction to ammonia): *nrfA*.

Archaea dominated this functional guild (80% of ammonia oxidisers), with all archaeal representatives belonging to the families Nitrosopumilaceae and Nitrososphaeraceae. This agrees with previous studies showing archaea to be the predominant drivers of ammonia oxidation on the planet (Leininger et al., 2006; Wuchter et al., 2006). The remaining 20% belong to the bacterial families Nitrospiraceae (phylum Nitrospirota) and Nitrosomonadaceae (phylum Pseudomonadota). Similarly, nitrite oxidation (nitrification) was phylogenetically constrained, occurring predominantly within Nitrospirota (Fig. 3), a pattern consistent with prior reports identifying this phylum as the most widespread nitrite oxidisers (Leung et al., 2022). In contrast, pathways like DNRA and denitrification (particularly nitrate, nitrite, and nitrous oxide reduction) were distributed across diverse bacterial and archaeal taxa, underscoring their functional versatility across lineages and environmental niches.

The taxonomic and metabolic insights generated here, from phylogenetically limited nitrifying taxa to more widespread denitrifiers, demonstrate *EcoFoldDB-annotate*’s capacity to disentangle complex microbial roles in biogeochemical cycling. This scalable and sensitive annotation framework thus allows efficient linking of microbial community structure to ecosystem-scale functions in large datasets.

### EcoFoldDB identifies microbial genomes encoding plant-microbe interaction traits

As a second use case example, we sought to identify potential plant growth-promoting members from the global soil microbiome. We used the EcoFoldDB annotations to identify soil MAGs encoding plant-microbe interaction traits. We found that many genomes encode multiple plant-microbe interaction traits, with 15.9% encoding two or more, and 2.5% encoding three or more (Fig. 4). A few of these traits were widespread, e.g., putrescine and spermidine production, which were present in most bacterial phyla. This is not surprising since biosynthesis of these polyamines is also prevalent in non-plant-associated bacteria given their diverse roles in bacterial physiology and growth, such as biofilm formation, motility, and siderophore production (Michael, 2018). The genetic capacity to produce the phytohormone auxin (indole-3-acetic acid, IAA) was also widespread, with 7.2% of MAGs containing at least one of the known IAA biosynthesis pathways (Fig. 4).

**Fig. 4.**
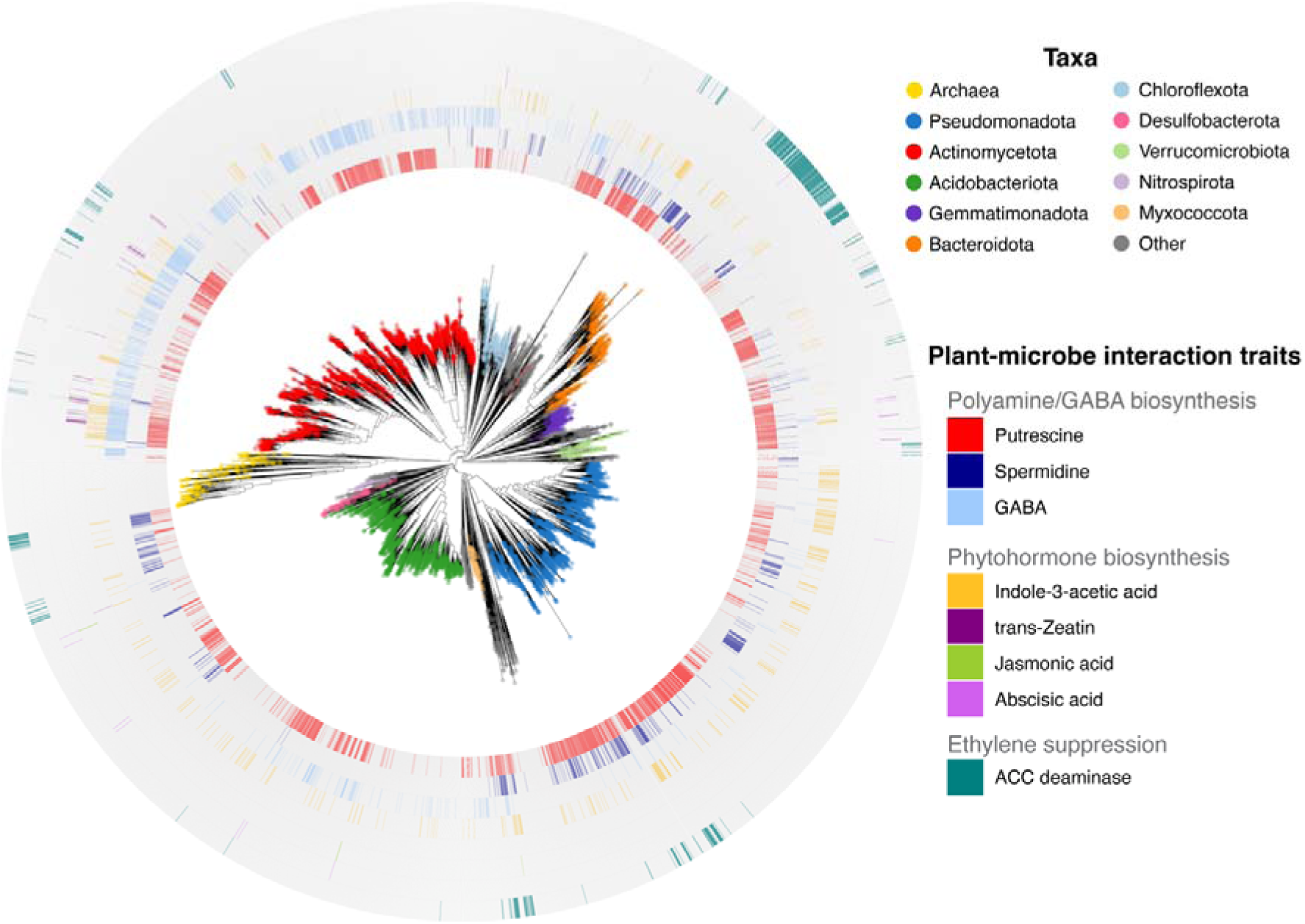
Plant-microbe interaction functions identified by *EcoFoldDB-annotate* among high-quality MAGs from the global soil microbiome. A whole genome phylogeny of high-quality bacterial and archaeal MAGs, derived from soil metagenomes across the globe. Coloured tip points on the tree indicate GTDB phyla, except for archaeal genomes, which have all been assigned a single domain-level colour (yellow tip points). Coloured rings indicate the genomes identified to have the genetic potential for plant-microbe interaction traits. As a minimum, the following genes were required for putrescine production: *speC* or (*speA* and *speB*) or (*speA* and *aguA* and *aguB*); spermidine production: (*speD*/*speH* and *speE*) or (*cansdh* and *nspC*); GABA (gamma-aminobutyric acid) production: *gadA*/*gadB* and *gadC*; indole-3-acetic acid (IAA) production via the indole-3-acetamide (IAM) pathway: *iaaM* or *iaaH*; IAA production via the indole-3-pyruvic acid (IPA) and tryptamine (TAM) pathways: *aldH*; IAA production via the indole-3-acetonitrile (IAN) pathway: *nit*; trans-Zeatin production: *log*; jasmonic acid production: *ehhadh* and *acox*; abscisic acid production: *aao*; brassinolide production: *br6ox* (not shown as no genomes were identified to carry this gene, Supplementary Table S2); gibberellin production: *ko* and *kao* (not shown as no genomes were identified to carry both genes, Supplementary Table S2); and ACC deaminase: *acdS*.

Other plant-microbe interaction traits, however, had strong taxonomic signals (Fig. 4). For example, 95% of trans-Zeatin producers and 76.2% of GABA producers belonged to the phylum Actinomycetota, largely from the order Mycobacteriales. ACC deaminase activity showed a strong Bacteroidota signature. Indeed, over half (55.2%) of all Bacteroidota MAGs encoded ACC deaminase, known to induce plant tolerance to abiotic stress (Glick et al., 2007). While ACC deaminase activity has been experimentally confirmed in Bacteroidota isolates (Maimaiti et al., 2007; Gandhi Pragash et al., 2009; Nadeem et al., 2009; Marques et al., 2010), research on this trait has historically focused on Pseudomonadota (Blaha et al., 2006; Nascimento et al., 2014). Thus, to our knowledge, the pronounced enrichment of ACC deaminase activity within Bacteroidota observed here has not been previously reported.

To identify possible novel plant growth promoting strains, we examined the subset of genomes encoding the greatest number of identified plant-microbe interaction traits, with 22 species-clustered MAGs being positive for four different traits (Fig. 5). Of these, 14 currently have no cultured species representative, with 10 lacking a cultured genus-level representative. The enrichment of encoded plant-microbe interaction traits, suggest that they likely have plant-growth promoting effects. Indeed, most of the species that do have a cultured representative have already been shown to comprise plant growth-promoting members (e.g., *Pseudomonas putida* (Hall et al., 1996; Ahemad and Khan, 2012; Costa-Gutierrez et al., 2020), *Pseudomonas sichuanensis* (a.k.a. *P. oryziphila*) (Yang et al., 2021), *Pseudomonas* sp024809425 (Izquierdo, 2017), *Pseudomonas furukawaii* (Rafique et al., 2022), *Rahnella variigena* (Raj et al., 2023)). The remaining species, which are also likely to comprise plant growth-promoting members, represent ideal candidates for targeted isolation and culturing. Such isolation efforts can be guided by genomic information from the MAG sequences, e.g., via optimisation of culture medium and conditions, including substrate utilisation, oxygen requirements, resistance to antibiotics, etc. (Pope et al., 2011; Karnachuk et al., 2021; Liu et al., 2022). This demonstrates how EcoFoldDB can help bridge ecological discovery and real-world application, translating metagenomic insights into actionable strategies for sustainable management.

**Fig. 5.**
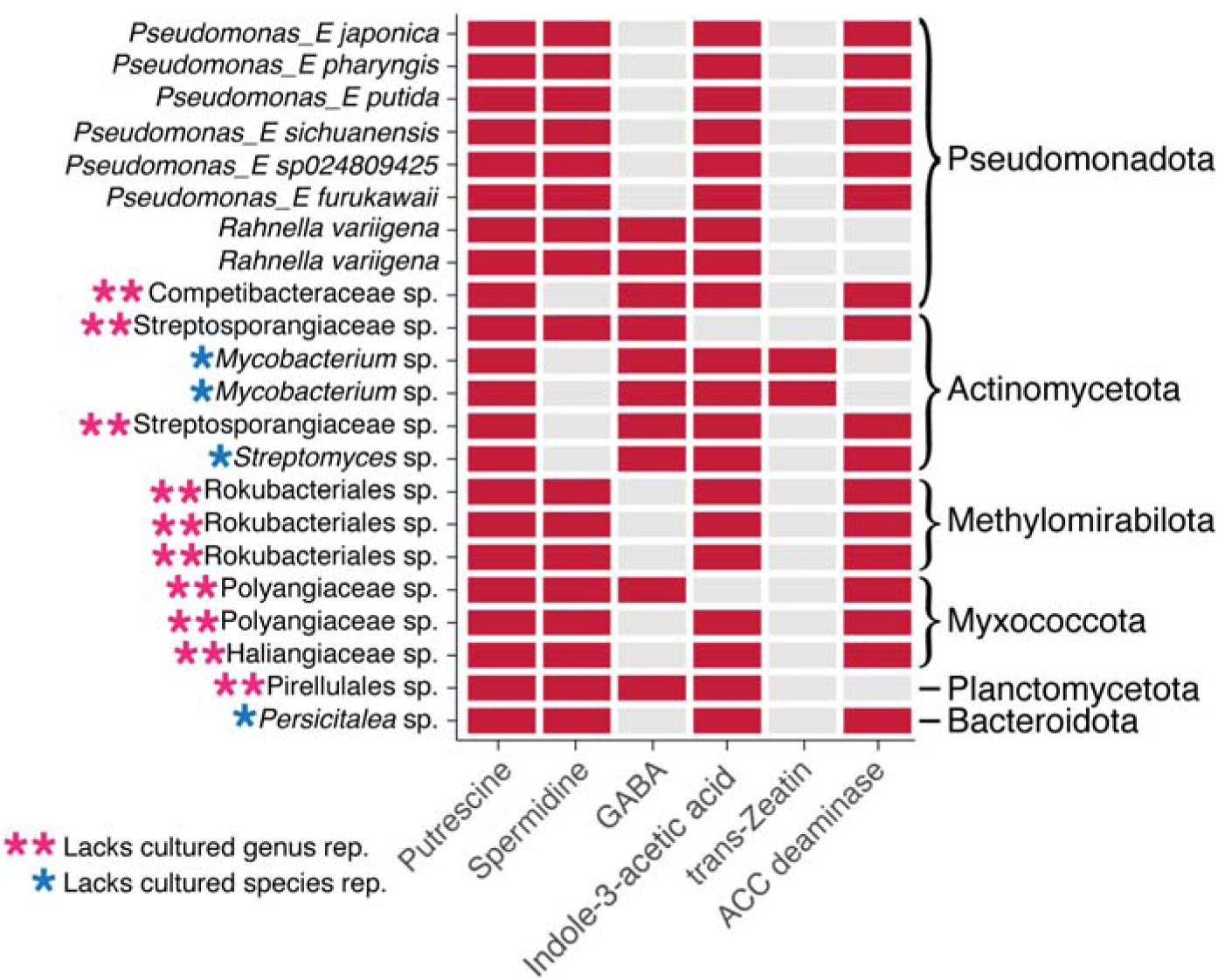
Identification of potential plant growth-promoting bacteria as candidates for targeted culturing. Soil MAGs encoding the greatest number of plant-microbe interaction traits (x-axis), as determined by *EcoFoldDB-annotate* are shown, labelled with their lowest named GTDB taxonomic rank (y-axis), and GTDB phylum (right). Each MAG represents a distinct 95% average nucleotide identity species cluster. The minimum gene sets for a genome to be considered positive (red rectangles) for each trait are as listed in Fig. 4. Strains lacking a cultured species representative are denoted with a single blue asterisk, and those also lacking a cultured genus representative are shown with double pink asterisks. Figure data, including the genome identifiers for these 22 MAGs, are available as Supplementary Table S3.

## Conclusion

Here, we introduce EcoFoldDB, a database of protein structures linked to ecologically relevant functions, and its companion pipeline, *EcoFoldDB-annotate*, for structure-guided functional profiling. *EcoFoldDB-annotate* achieves highly sensitive and accurate annotations, scalable to metagenomic datasets (millions of proteins). We showcase the use of *EcoFoldDB-annotate* on a global soil microbiome MAG catalogue, revealing key insights into the taxonomic distribution of important nitrogen cycling pathways, and plant-microbe interaction traits, advancing our understanding of important ecosystem processes. We anticipate that EcoFoldDB will serve as a valuable resource for microbial ecology studies, enabling researchers to extract ecological insights from environmental genomes and metagenomes, and accelerating discoveries in microbiome science.

## Availability of data and material

EcoFoldDB and *EcoFoldDB-annotate* pipeline are available via https://github.com/timghaly/EcoFoldDB. Genomic and protein sequences for the high-quality soil MAGs are available via https://doi.org/10.5281/zenodo.15054612. EcoFoldDB annotations of these genomes are available as Supplementary Table S2.

## Competing interests

The authors declare that they have no competing interests.

## Funding

This work was funded by a Macquarie University Research Fellowship (T.M.G) and the ARC Centre of Excellence in Biology (S.G.T).

## Authors’ contributions

T.M.G was involved in the project conception and design, conducted the research, and wrote the manuscript draft. V.R., D.R and E.C were involved in pipeline testing and analysis. S.G.T was involved in project design and management. All authors contributed to the final editing of the manuscript.

## Acknowledgements

We would like to thank the developers of all the software used in this study, particularly Foldseek and ProstT5, without which, this work would not have been possible. TMG would also like to thank Mary, Saoirse, and Maebh Ghaly for their loving support.

## Notes

### Competing Interest Statement

The authors have declared no competing interest.

### Summary of Updates

Methods updated to improve benchmarking dataset. This has affected benchmarking results (Fig. 2). EcoFoldDB has been updated to version 2.0. Figs. 3-5 have and Supplementary Tables S1-3 have been updated accordingly.

